# Sleep-Related Respiratory Disruption is Associated with Altered Spindle Morphology and Poorer Attention in Children

**DOI:** 10.64898/2026.07.06.736760

**Authors:** Ido Haber, Tamara P. Taporoski, Beth Peterson, Camilla Matthews, Tony Kille, Annika Myers, Brady Riedner, Emma Strainis, Ana Maria Vascan, Stephanie G. Jones

## Abstract

**Study Objectives:** To determine whether sleep-related respiratory disruption is associated with regionally specific alterations in sleep spindle topography and whether hypopnea-sensitive spindle features are associated with attentional performance in children.

**Methods:** We recorded overnight high-density EEG in children across a wide range of respiratory disruption severity. Slow and fast spindle metrics were extracted per channel, and channel-wise regression models characterized topographic associations with hypopnea index (HI). Cluster-based permutation testing controlled for multiple comparisons. Hierarchically defined regions of interest were tested as predictors of attentional performance on the Test of Variables of Attention (TOVA).

**Results:** Canonical slow-anterior and fast-posterior spindle organization was detectable across the cohort. Two HI-related topographic effects survived cluster-based permutation correction: higher HI was associated with shortened anterior fast spindle duration and with slower anterior slow spindle peak frequency. In cognitive models, anterior fast spindle duration was the strongest and most consistent predictor of attentional performance, associated with higher signal detection sensitivity, fewer omission errors, and fewer commission errors. By contrast, slow spindle peak frequency showed no attentional associations.

**Conclusions:** Pediatric respiratory disruption is associated with regionally specific alterations in spindle morphology rather than global spindle reduction. Shortened anterior fast spindle duration showed convergent respiratory and attentional associations, suggesting that localized spindle integrity may provide a neurophysiological marker of cognitive vulnerability in pediatric sleep-disordered breathing beyond conventional clinical respiratory metrics.

**Highlights:**

- Higher hypopnea index was associated with shortened anterior fast spindle duration and slower anterior slow spindle peak frequency
- Anterior fast spindle duration predicted attentional performance in children
- Respiratory disruption may impair attention by disrupting thalamocortical spindle activity

**Statement of Significance:** Children with sleep-disordered breathing can show neurocognitive difficulties that are not well explained by standard respiratory indices alone. This study uses high-density EEG to show that hypopnea burden is associated with regionally specific alterations in sleep spindle morphology, particularly shortened anterior fast spindle duration, rather than a global reduction in spindle occurrence. Anterior fast spindle duration was also the spindle feature most consistently associated with attentional performance. These findings suggest that localized spindle morphology may provide a sleep-neurophysiological readout of cognitive vulnerability in pediatric sleep-disordered breathing. Longitudinal and treatment studies are needed to determine whether spindle duration changes with respiratory improvement and whether such changes track cognitive recovery.

## Introduction

Sleep-disordered breathing (SDB) represents a spectrum of respiratory disturbances during sleep that affects an estimated 1–5% of children in the general population, with prevalence rates substantially higher among children referred for clinical evaluation of snoring, adenotonsillar hypertrophy, or obesity [1]. Even mild forms of SDB, including primary snoring and upper airway resistance syndrome, are increasingly recognized as clinically meaningful given their associations with neurocognitive impairment, behavioral dysregulation, and diminished academic performance [2,3,4,5]. The hypopnea index (HI), which quantifies partial airway obstruction events per hour of sleep, captures a component of respiratory disruption that may be particularly relevant to sleep microarchitecture, as hypopneas produce intermittent hypoxemia and arousal-related fragmentation without necessarily meeting the full criteria for obstructive apnea [1].

Sleep spindles are transient thalamocortical oscillations generated by interactions between the thalamic reticular nucleus and thalamocortical relay neurons, appearing as waxing-and-waning bursts of 10–16 Hz activity during non-rapid eye movement (NREM) sleep [6,7]. A fundamental distinction exists between slow spindles (approximately 10–12 Hz), which predominate over frontal cortical regions, and fast spindles (approximately 12–16 Hz), which are maximally expressed over centroparietal areas [8,9,10]. This anterior-posterior dissociation is thought to reflect at least partially separable thalamocortical circuits [6,8]; although the functional significance of the two spindle types remains incompletely understood, fast spindles in particular couple with the hippocampal-cortical dialogue that supports memory consolidation [6]. How this topographic organization matures across childhood has been characterized primarily with conventional or comparatively lower-density EEG montages [11,12]; the high-density electroencephalography (hdEEG) used here offers the spatial resolution to resolve finer regional variation that lower-density systems may not capture.

A growing body of evidence indicates that SDB and obstructive sleep apnea (OSA) alter sleep spindle characteristics, though the majority of this work has been conducted in adult populations [13]. Large-scale characterization studies have established that spindle features — including density, amplitude, duration, and frequency — represent distinct, heritable dimensions of sleep microarchitecture with topographic specificity [14]. The few pediatric investigations that exist suggest that OSA can alter spindle activity [15,16] and broader sleep microstructure [17], but they have generally relied on limited-montage polysomnography, restricting the ability to characterize topographic alterations. Whether the canonical slow-anterior and fast-posterior spindle organization is disrupted by respiratory disturbance in children, and whether such disruption exhibits regional selectivity, remains an open question. This gap is particularly notable given evidence that childhood represents a critical period for thalamocortical circuit maturation, during which spindle characteristics undergo marked developmental changes [18,19,20,21].

Sleep spindles have been consistently linked to cognitive functioning, with particular relevance to memory consolidation and broader cognitive performance [22,23]. In children, spindle activity has been associated with neurocognitive performance, general cognitive abilities, and learning efficiency [21,24,25]. Of particular interest for the present investigation, emerging evidence suggests that the multidimensional nature of spindle morphology — including duration, amplitude, and frequency — may capture aspects of thalamocortical function not reflected by spindle occurrence rate alone [14,26]. The Test of Variables of Attention (TOVA), a continuous performance test that yields measures of signal detection sensitivity, sustained attention, inhibitory control, and processing speed, provides a well-validated assessment of sustained attention and response control in children [27,28].

Prior pediatric work has shown that children with OSA can exhibit reduced spindle activity relative to controls [15], and that children with SDB and daytime sleepiness show lower spindle activity than children with SDB without sleepiness [16]. These findings support sleep spindles as a clinically relevant dimension of pediatric SDB microstructure, but leave unresolved whether respiratory disruption affects spindles as a global reduction in occurrence or as a regionally selective disturbance in specific spindle features. This distinction is important because spindle features are not interchangeable markers of thalamocortical function [14,26]. A child may show preserved spindle occurrence but altered spindle stability, or altered oscillatory frequency without parallel cognitive relevance [14,26]. High-density EEG allows these dimensions to be separated topographically and tested against cognitive outcomes.

The present study leverages hdEEG to characterize the topographic distribution of sleep spindle activity in children and to examine how the severity of respiratory disruption, indexed by HI, relates to spindle topography and attentional performance. We pursue three aims: (1) to characterize whether canonical slow-anterior and fast-posterior spindle organization is detectable in this pediatric SDB-enriched cohort, (2) to determine whether higher HI is associated with regional shifts in spindle expression and alterations in spindle morphology, and (3) to evaluate whether spindle features identified as sensitive to respiratory disruption are independently associated with attentional performance. We hypothesized that HI would be associated with altered spindle activity, based on prior pediatric evidence that children with OSA show reduced spindle activity relative to controls [15]. Because spindle features vary by topography and represent separable dimensions of sleep microarchitecture [8,14,26], we expected these HI-related effects to be regionally and morphologically specific rather than diffuse. Finally, given pediatric evidence linking spindle activity to cognitive performance [21,24,25], we hypothesized that HI-sensitive spindle features would be associated with attentional performance.

## Methods

### Participants

Participants were drawn from a single-site observational study of pediatric sleep neurophysiology spanning the full range of sleep-disordered breathing. Children were recruited both from clinical referrals for suspected obstructive sleep apnea (e.g., otolaryngology or adenotonsillectomy evaluation) and from the surrounding community through advertising and word of mouth, capturing children with minimal symptoms through polysomnography-defined disease. The study was approved by the University of Wisconsin–Madison Institutional Review Board (protocol 2017-0681); a parent or legal guardian provided written informed consent and children provided age-appropriate assent before any study procedures. Recruitment spanned 2018 through early 2020 and was halted by the COVID-19–related laboratory closure; 80 children provided consent and 72 completed overnight study procedures before the interruption. Eligibility required an age within the preadolescent study range, a caregiver able to provide consent and developmental and medical history, English-language ability sufficient to complete the study tasks, and tolerance of the overnight EEG and polysomnography setup. Children were excluded for a history of significant head trauma, unstable medical illness, current psychotropic medication use, or intellectual disability or substantial cognitive impairment. Among children who completed the overnight recording, 10 were excluded because technical problems with the hdEEG or polysomnographic sleep recording precluded reliable spindle analysis, yielding a final spindle analytic sample of 62 children (33 boys, 29 girls; mean age = 7.7 years, range: 3.0–11.0). Of these, 57 had complete TOVA data; one was excluded as an extreme outlier (z > 3 on log-transformed reaction time), yielding a cognitive analytical sample of *N* = 56.

### Instruments

Overnight hdEEG was acquired with a 256-channel HydroCel Geodesic Sensor Net (Electrical Geodesics Inc., Eugene, OR) [29] run on Compumedics Neuvo amplifiers, with gel-filled electrodes referenced to the vertex during acquisition and digitized at 500 Hz. Clinical polysomnography was recorded concurrently and included electrooculography, chin and leg electromyography, pulse oximetry, respiratory effort belts, and airflow sensors. Sleep was staged visually in 30-second epochs and respiratory events were scored according to American Academy of Sleep Medicine criteria, with clinical interpretations reviewed by a board-certified sleep medicine physician. Respiratory indices were expressed as events per hour of sleep: the hypopnea index (HI) as hypopneas per hour and the apnea–hypopnea index (AHI) as apneas plus hypopneas per hour. Because both clinically referred and community-recruited children spanned a continuous range of objectively scored respiratory burden, HI was modeled as a continuous exposure rather than by recruitment source or diagnostic category.

To assess attentional performance, participants completed the Test of Variables of Attention (TOVA), a computerized fixed-interval continuous performance test [28]. The TOVA is a 21.6-minute task that measures sustained attention, response speed, and inhibitory control. During the assessment, participants are presented with simple geometric stimuli and are instructed to press a button as quickly as possible when the target appears and to withhold their response when a non-target appears.

The assessment is divided into four equal segments, or quarters, of 5.4 minutes each. The present analyses focused on the first quarter (Q1), which consists of 162 stimuli presented at a 2-second inter-stimulus interval and represents the initial low-target-frequency phase of the task. Primary outcome variables included d-prime, a signal-detection metric indexing participants’ discrimination of targets from non-target stimuli; mean reaction time (in milliseconds), indexing processing speed; omission errors, reflecting missed target responses; and commission errors, reflecting responses to non-target stimuli.

### Study Protocol

Testing was conducted across an initial daytime assessment visit and a subsequent overnight laboratory visit; the present analyses use the baseline overnight recording and the subsequent morning TOVA performance. On the overnight visit, children arrived approximately 2 hours before their habitual bedtime to allow hdEEG net fitting and polysomnography sensor placement, and a caregiver was permitted to remain overnight. Children completed evening waking procedures before all-night polysomnography and hdEEG recording under ad libitum sleep, after which morning neurobehavioral testing, including the TOVA, was administered.

### EEG Preprocessing and Spindle Detection

hdEEG data were processed offline in MATLAB using EEGLAB [30] and custom laboratory scripts. After importing electrode locations and sleep and arousal annotations and removing non-EEG channels, data were low-pass filtered at 40 Hz, resampled to 200 Hz, and high-pass filtered at 0.5 Hz. To eliminate low-frequency line-noise sub-harmonics without distorting the underlying broadband EEG signal or phase-locking characteristics within the σ band, narrowband line-related noise at 10, 20, and 30 Hz was attenuated using Zapline-plus [31]. Channels with poor signal for more than 30% of the night were removed, and face and neck electrodes were excluded from scalp topographic analyses; all spindle metrics and topographic analyses used a standardized 172-channel montage covering the scalp surface (Figure S1). Removed channels were reconstructed by spherical-spline interpolation when overall data quality remained adequate, and the data were re-referenced to the average of the retained scalp channels.

Spindle detection was restricted to artifact-free NREM sleep (stages N2 and N3), which constituted the analysis window (Figure 1a). Spindles were detected using YASA, an open-source Python package [32], applied independently to each of the 172 EEG channels. The detection procedure (Figure 1b) involved bandpass filtering the EEG signal in the σ frequency range (10–16 Hz), computing the analytic amplitude envelope via the Hilbert transform, and identifying threshold-crossing events. Detected events were required to satisfy the following criteria: duration between 0.5 and 2.0 seconds, minimum inter-spindle interval of 500 milliseconds, relative σ power ≥ 0.2, correlation with a σ-frequency template ≥ 0.65, and a moving root-mean-square amplitude exceeding its mean by at least 1.5 standard deviations. Each detected spindle was classified as slow (10 to <12 Hz) or fast (12 to 16 Hz) based on its peak oscillation frequency [8]. For each channel and spindle type, four metrics were extracted: total count, mean amplitude (microvolts), mean duration (seconds), and mean peak frequency (Hz). Spindle density (events per minute) was computed by dividing the total spindle count by the duration of artifact-free N2 + N3 sleep. Channels with zero detected events were assigned a spindle density of zero. For the morphological metrics (amplitude, duration, and peak frequency), these channels were treated as missing data, because morphological estimation requires the presence of at least one detected event.

**Figure 1.**
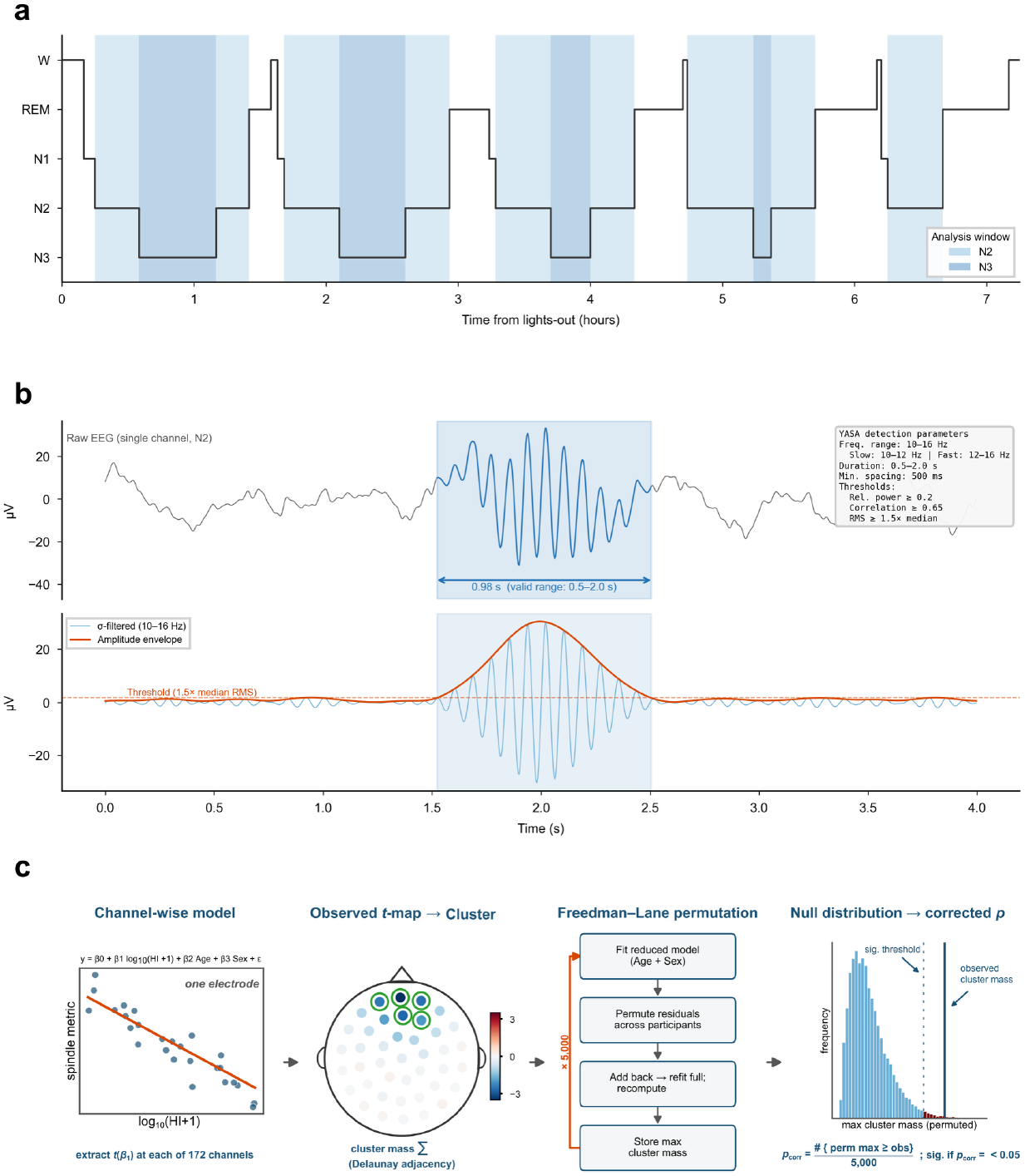
Sleep architecture, spindle detection, and statistical methodology. (a) Representative pediatric sleep hypnogram illustrating the analysis window. Spindle detection was restricted to NREM stages N2 (light shading) and N3 (dark shading). (b) Illustration of the YASA spindle detection procedure applied to a single EEG channel during N2 sleep. Upper trace: raw EEG signal with a detected spindle event highlighted (blue shading) and duration annotation. Lower trace: σ-band filtered signal (10–16 Hz) with amplitude envelope (orange) and detection threshold (dashed line; moving root-mean-square amplitude exceeding its mean by 1.5 standard deviations). Detection parameters are summarized in the inset. (c) Cluster-based permutation testing pipeline (left to right): a channel-wise regression of each spindle metric on HI (adjusted for age and sex) yields an HI regression coefficient and t-value at each of the 172 channels; the observed t-map is thresholded (|t| > 2.0) and grouped into spatially contiguous clusters via Delaunay-triangulation adjacency, with each cluster summarized by its mass (sum of |t|); the null distribution is generated by Freedman–Lane residual permutation across participants (5,000 iterations, a single relabeling applied across all channels per iteration to preserve spatial covariance); and the corrected p-value for each observed cluster is the proportion of permutation maxima greater than or equal to its mass.

### Statistical Analysis

Topographic associations between HI and spindle metrics were examined using channel-wise linear regression models. For each of the 172 channels, each spindle metric was modeled as:

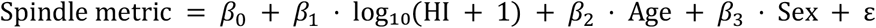

Age was mean-centered, and HI was transformed as log_10_(HI + 1) and mean-centered. The log transformation reduced right skewness while retaining children with zero or near-zero respiratory events. The regression coefficient β_1_ (the slope per unit log_10_(HI + 1), in each spindle metric’s native units) and its associated *t*-statistic were extracted at each channel to characterize the direction, magnitude, and significance of associations between HI and spindle metrics across the scalp. Channels reaching significance at *p* < 0.05 (uncorrected) were identified to characterize the spatial distribution of effects, and spatially coherent clusters were defined for subsequent region-of-interest (ROI) analyses. These uncorrected topographic patterns were then evaluated using cluster-based permutation testing to determine which effects survived correction for multiple comparisons.

ROIs were hierarchically classified as confirmatory or exploratory according to whether the underlying HI effect survived permutation-based correction. ROI channel membership was defined in one of two ways: the anterior slow spindle peak-frequency ROI was taken directly as its permutation-corrected cluster, whereas the remaining ROIs — including the cluster-corrected anterior fast spindle duration effect — were defined from spatially coherent uncorrected channel-wise effects (*p* < 0.05, with adjacent trending channels at *p* < 0.08 retained to capture the full spatial extent of each effect). The two exploratory ROIs (posterior slow amplitude, anterior slow duration) were retained only to assess whether behavioral associations were specific to corrected effects or generalized across spindle features. For each participant, a single representative value was computed for each ROI by averaging the relevant spindle metric across all channels within the ROI.

Cluster-based permutation testing [33] was used to evaluate whether the observed topographic associations exceeded what would be expected by chance while accounting for spatial dependence among neighboring electrodes (Figure 1c). The test statistic at each electrode was the *t*-value for the HI coefficient from the channel-wise multiple regression described above (spindle metric on HI, age, and sex). Electrodes exceeding a cluster-forming threshold of |*t*| > 2.0 were grouped into spatially contiguous clusters using Delaunay triangulation-based channel adjacency (MNE-Python) [34], and the cluster-level statistic was the sum of |*t*|-values within each cluster (cluster mass). The null distribution was generated by Freedman-Lane residual permutation [35], which preserves the covariate adjustment under the null: the reduced model (age and sex, omitting HI) was fit, its residuals were permuted across participants — using a single relabeling applied to every electrode within each permutation to preserve the spatial covariance structure — added back to the reduced fitted values, and the full model was refit to recompute the HI *t*-map. The maximum cluster mass was retained on each of 5,000 permutations, and the corrected *p*-value for each observed cluster was the proportion of permutation maxima greater than or equal to its mass. Clusters with corrected *p* < 0.05 were considered statistically significant.

As a follow-up sensitivity analysis prompted by visual age-stratified maps, we also tested whether age moderated HI-spindle associations. For each channel and metric, we fit a model including age, HI, their interaction (HI × Age), and sex. This sensitivity analysis was used to evaluate whether respiratory burden affected spindle topography differently by developmental stage.

Associations between spindle ROI metrics and attentional performance were examined using linear regression models. Quarter 1 TOVA variables (d-prime, omission errors, commission errors, and mean reaction time) served as dependent variables, modeled separately as a function of each ROI metric:

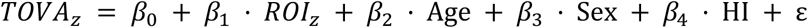

All models were adjusted for age, sex, and HI. To meet assumptions of normality, omission and commission errors were square-root transformed to account for zero-inflation, and mean reaction time was log-transformed to correct for left-tail skewness. One participant with a standardized residual exceeding 3 standard deviations on reaction time was excluded, yielding a cognitive analytical sample (ROI and TOVA data present) of *N* = 56. All dependent variables and ROI predictors were standardized (z-scored) to facilitate comparison of effect sizes across models. We used the behavioral models strictly to confirm the topographic findings rather than as an independent exploratory analysis. Accordingly, we focused interpretation only on spindle features that met two criteria: they survived cluster-based correction in the topographic analysis, and they were robustly associated with multiple TOVA outcomes.

Analyses were implemented in MATLAB and Python. EEG preprocessing was performed in MATLAB (R2023b; MathWorks) with EEGLAB (v2023.1) and the Zapline-plus extension (v1.2.1); spindle detection used YASA (v0.6.5), topographic cluster-based permutation testing used MNE-Python (v1.11.0), and channel-wise and behavioral regression models were fit in Python using statsmodels [36] and SciPy [37]. Statistical tests were two-sided with α = .05 unless otherwise specified.

## Results

### Sample Characteristics

The final sample included 62 children (33 boys, 29 girls; mean age = 7.7 years, range: 3.0–11.0) with a wide range of HI values (mean HI = 3.1, range: 0.1–34.1; mean AHI = 4.7, range: 0.5–36.2). Demographic characteristics and descriptive statistics for spindle metrics and cognitive performance are summarized in Table 1. Of these, 56 participants with complete TOVA data and no extreme outliers on transformed variables constituted the cognitive analytical sample (ROI and TOVA data present).

**Table 1.** Demographic characteristics, sleep spindle metrics, and attentional performance.

|  | Value | *n* |
| --- | --- | --- |
| **Demographics** |  |  |
| Age (years) | 7.7 (3.0, 11.0) | 62 |
| Boys / Girls | 33 (53%) / 29 (47%) | 62 |
| **Respiratory indices** |  |  |
| Hypopnea index (events/hr) | 3.1 (0.1, 34.1) | 62 |
| Apnea–hypopnea index (events/hr) | 4.7 (0.5, 36.2) | 62 |
| **Spindle count (total events)** |  |  |
| Slow (10–12 Hz) | 189.0 (1.6, 699.7) | 62 |
| Fast (12–16 Hz) | 130.6 (0.2, 628.0) | 62 |
| **Spindle density (events/min)** |  |  |
| Slow (10–12 Hz) | 0.6 (0.0, 2.0) | 62 |
| Fast (12–16 Hz) | 0.4 (0.0, 1.9) | 62 |
| **Spindle amplitude (μV)** |  |  |
| Slow (10–12 Hz) | 55.3 (33.8, 94.0) | 62 |
| Fast (12–16 Hz) | 54.8 (27.6, 79.8) | 62 |
| **Spindle duration (s)** |  |  |
| Slow (10–12 Hz) | 0.9 (0.7, 1.0) | 62 |
| Fast (12–16 Hz) | 0.8 (0.6, 1.0) | 62 |
| **Attentional performance (TOVA Q1)** |  |  |
| d-prime | 4.6 (0.5, 8.5) | 56 |
| Omission errors | 9.5 (0.0, 52.2) | 56 |
| Commission errors | 3.2 (0.0, 29.5) | 56 |
| Mean reaction time (ms) | 607.9 (353.0, 838.4) | 56 |

*Note.* Continuous variables presented as mean (min, max); categorical variables as *n* (%). Spindle metrics represent channel-averaged values across 172 electrodes. TOVA = Test of Variables of Attention, Quarter 1.

Topographical analysis of spindle metrics was supported by high channel-wise data integrity: on average, 99% of participants contributed valid slow spindle metrics and 96% contributed valid fast spindle metrics per channel, with a minimum of 54 valid entries (of 62) at every channel.

### Normative Topography of Slow and Fast Spindles

We first characterized the spatial distribution of spindle activity across the full sample (Figure 2a–d).

**Figure 2.**
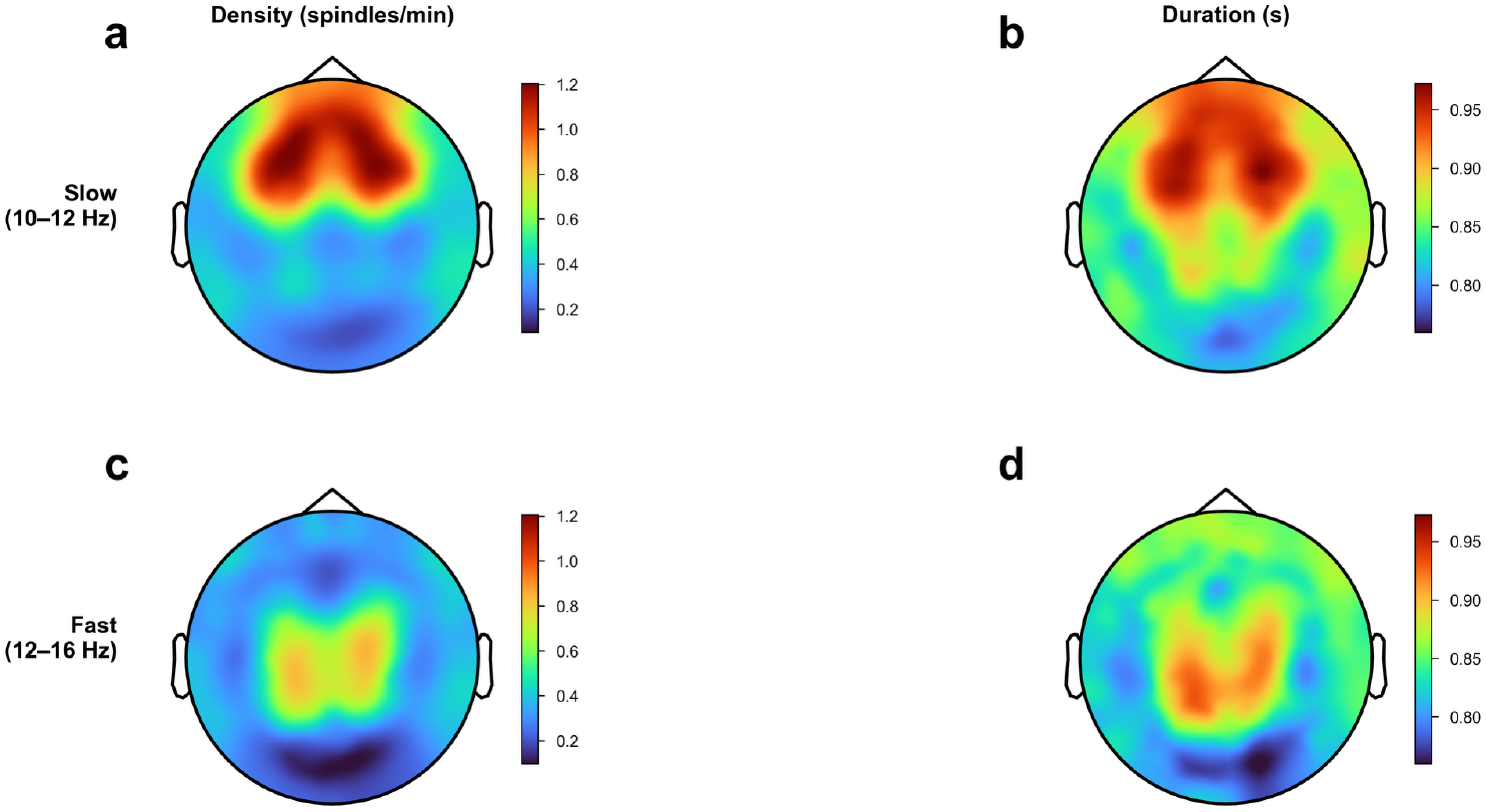
Normative topography of slow and fast sleep spindles. Panels a–b show slow spindle density and duration (seconds), averaged across participants. Panels c–d show corresponding metrics for fast spindles. Color scales are matched within each metric across bands to facilitate direct comparison. All maps are displayed in standard electrode space following interpolation across 172 channels.

Slow spindles (10–12 Hz) exhibited a robust anterior-frontocentral predominance (Figure 2a–b). Peak spindle density was localized over frontal electrodes (∼350 total spindles; ∼0.6 events/min), while amplitude showed a broad midline-frontal peak (∼70 microvolts). Duration was spatially homogeneous, ranging between 0.8 and 0.9 seconds with a slight frontocentral maximum.

Fast spindles (12–16 Hz) demonstrated a centroparietal predominance (Figure 2c–d). Peak production was localized to centroparietal regions (∼250 total spindles; ∼0.4 events/min), with a midline-central amplitude peak (∼70 microvolts) and a characteristic central-posterior power increase. Fast spindle duration largely paralleled density across the scalp: the longest durations (∼0.91 s) coincided with the centroparietal regions of peak spindle production, whereas the shortest durations (∼0.78 s) occurred over lower-density posterior and peripheral channels. These spatial patterns are consistent with canonical slow and fast spindle topographies described in both adult and pediatric literature, indicating that the fundamental slow-anterior/fast-posterior organization was detectable in this sample.

### Associations Between Hypopnea Index and Spindle Topography

We next examined associations between HI and spindle characteristics using channel-wise linear regression models adjusted for age and sex. HI was not associated with a diffuse reduction in spindle production. Neither slow nor fast spindle count/density showed cluster-corrected associations with HI, indicating that respiratory disruption was not primarily expressed as loss of spindle occurrence. Instead, the strongest effects emerged in spindle morphology, particularly anterior fast spindle duration. Two effects survived cluster-based permutation correction and are presented in Figure 3, followed by exploratory uncorrected topographic patterns retained for comparison and displayed in Figure 4.

**Figure 3.**
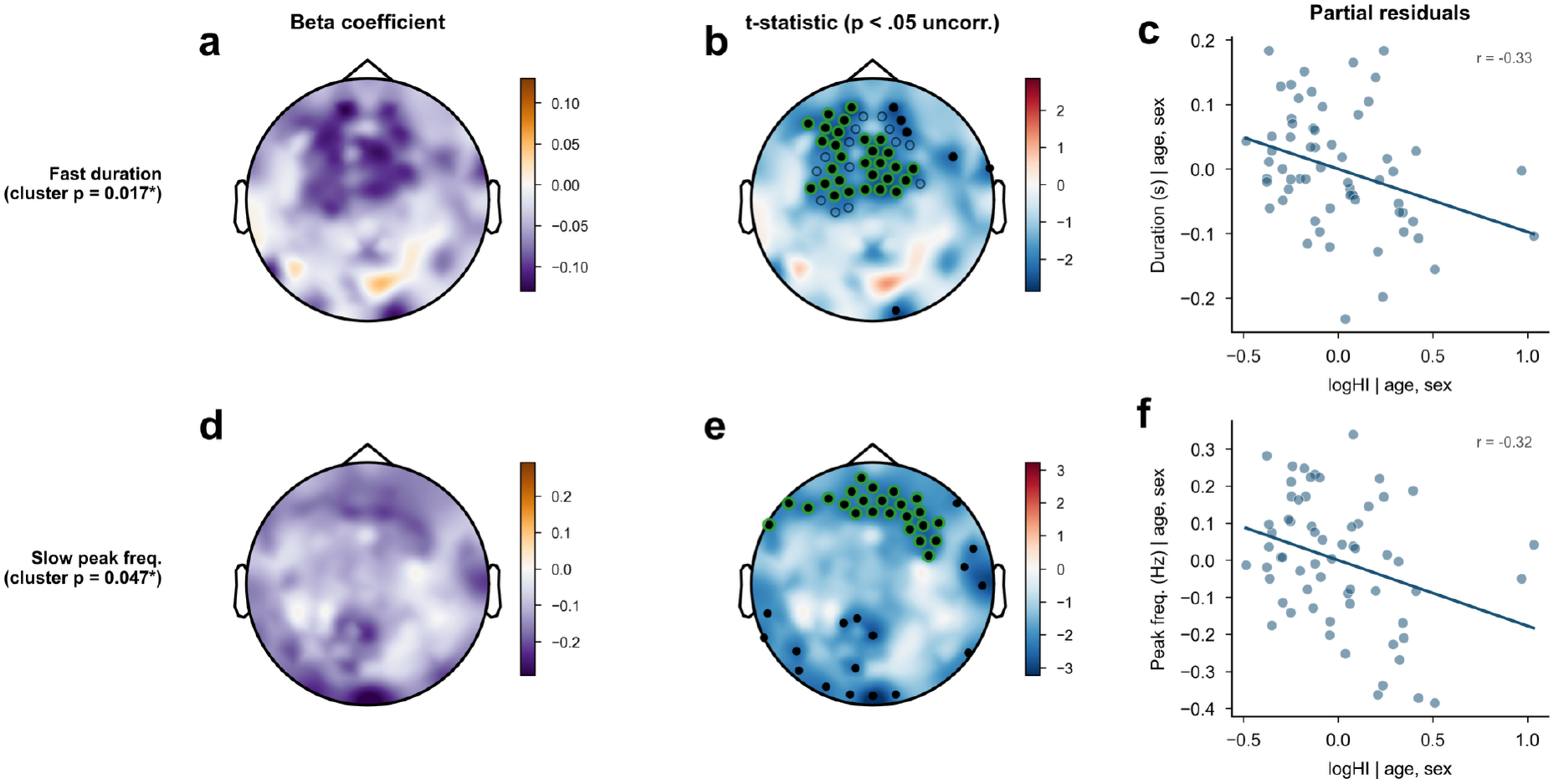
Cluster-corrected associations between hypopnea index and spindle metrics. Two topographic effects survived cluster-based permutation correction. Top row (a–c): fast spindle duration showed a large anterior cluster of shortened duration with increasing HI (corrected p = 0.017). Bottom row (d–f): slow spindle peak frequency showed an anterior cluster of slower oscillation frequency with increasing HI (corrected p = 0.047). Left column: regression coefficients (raw metric units; warm = positive, cool = negative). Center column: t-statistics. Filled black circles mark every channel significant at p < 0.05 uncorrected, including spatially isolated channels that do not enter any ROI or surviving cluster and are shown only to depict the full uncorrected map. Green open circles outline the cluster that survived permutation correction (fast duration, 29 channels; slow peak frequency, 24 channels); in the top row, black open circles additionally outline the 47-channel anterior ROI averaged for the behavioral analyses, whereas for slow peak frequency the surviving cluster is itself the behavioral ROI. Right column: partial-residual scatter plots (adjusted for age and sex) of ROI-averaged metric versus HI with linear regression fit.

**Figure 4.**
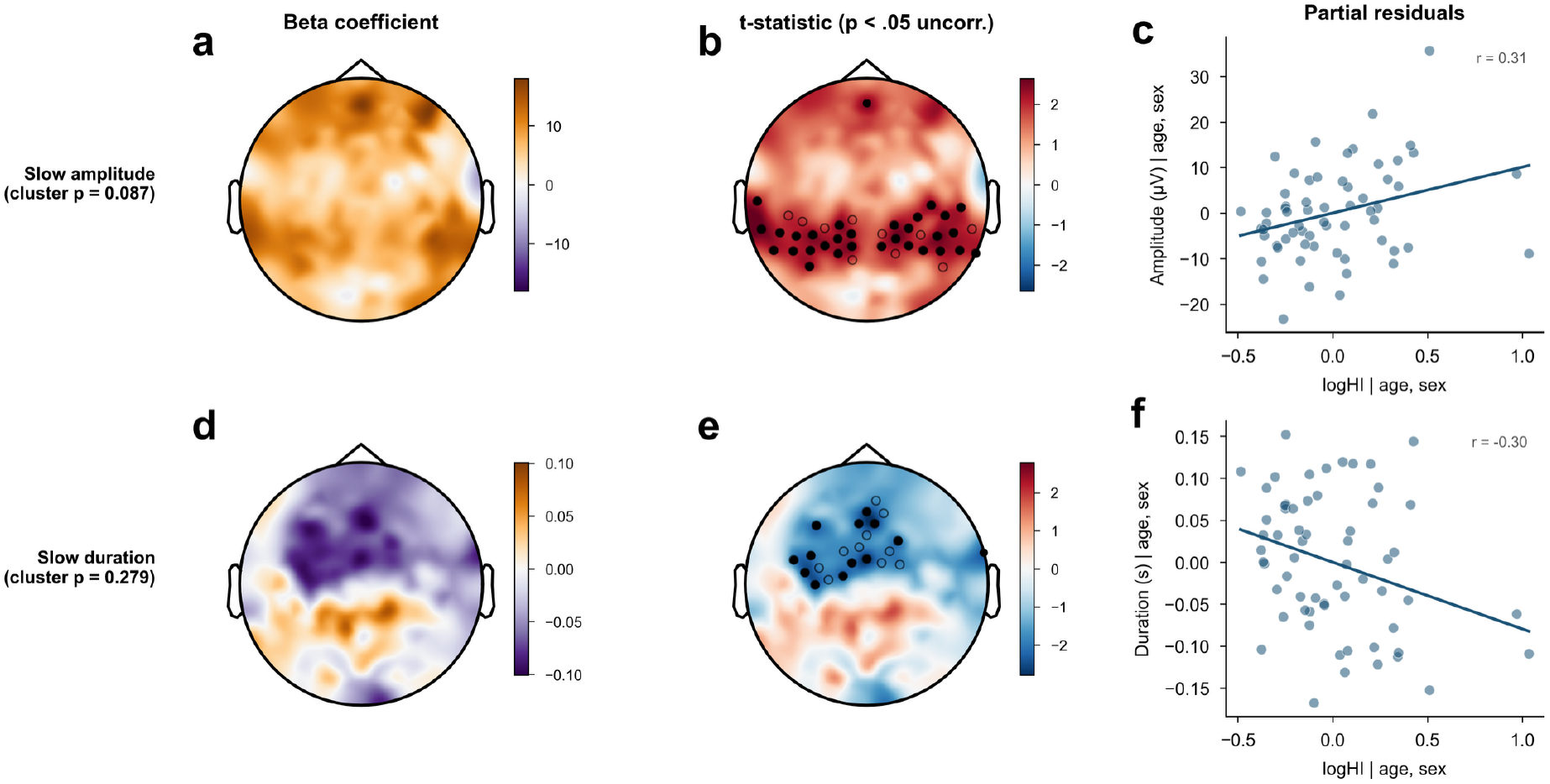
Exploratory uncorrected topographic associations between hypopnea index and spindle metrics retained for comparison with cluster-corrected effects. Top row (a–c): posterior slow spindle amplitude showed positive associations with HI. Bottom row (d– f): anterior slow spindle duration showed negative associations with HI. Neither effect survived cluster-based permutation correction (slow amplitude cluster p = 0.087; slow duration cluster p = 0.279). Left column: regression coefficients (raw metric units). Center column: t-statistics. Filled black circles mark every channel significant at p < 0.05 uncorrected, including spatially isolated channels that do not enter the ROI; open black circles outline the exploratory ROI averaged for the behavioral analyses (posterior slow amplitude, 40 channels; anterior slow duration, 22 channels). Right column: partial-residual scatter plots (adjusted for age and sex) of ROI-averaged metric versus HI with linear regression fit. These exploratory clusters were retained to evaluate whether behavioral associations were specific to corrected effects or generalized across spindle features.

### Cluster-Corrected Associations

Fast spindle duration demonstrated the most pronounced and widespread topographic effect in the analysis (Figure 3a–c): a large anterior cluster of 35 channels showed significant negative associations with HI at the uncorrected level, with a 29-channel subset surviving cluster-based permutation correction (corrected *p* = 0.017). Higher respiratory disruption was associated with shorter fast spindle durations in anterior regions. Individual-level data confirmed a consistent negative association between HI and ROI-averaged fast spindle duration (Figure 3c). This effect was robust to leave-one-subject-out reanalysis and subject-level bootstrap resampling (Figure S4). Fast spindle count, density (Figure S5), and amplitude did not exhibit robust topographical associations with HI.

Slow spindle peak frequency — the mean oscillation frequency of detected slow spindles within each channel — also demonstrated a cluster-corrected association with HI (Figure 3d–f). Channel-wise regressions revealed widespread negative associations: 41 of 172 channels showed significant effects at *p* < 0.05 (uncorrected), all in the negative direction. Cluster-based permutation testing confirmed a significant 24-channel anterior cluster (corrected *p* = 0.047), indicating that higher respiratory disruption is associated with slower oscillation frequencies within the slow spindle band. Individual-level data showed consistent negative relationships between HI and ROI-averaged slow spindle peak frequency (Figure 3f).

### Uncorrected Topographic Patterns

Beyond the cluster-corrected effects, additional topographic patterns were observed that, while not surviving correction for multiple comparisons, showed spatially coherent associations with HI (Figure 4). Slow spindle amplitude showed significant positive associations with HI across a posterior region of 31 channels (uncorrected, *p* < 0.05; Figure 4a–c), which together with contiguous trending channels (*p* < 0.08) formed the 40-channel posterior ROI averaged for the behavioral analysis, indicating that higher respiratory disruption is linked to greater posterior slow spindle amplitude. Individual-level data showed a positive trend between HI and slow spindle amplitude within the posterior ROI (Figure 4c). Slow spindle duration showed significant negative associations with HI across 13 anterior channels (uncorrected, *p* < 0.05; Figure 4d–f), extended to a 22-channel ROI by the same criterion, with higher HI predicting shorter spindle durations. Slow spindle density showed a spatial trend toward posterior increases with HI, but this pattern did not yield a spatially coherent cluster meeting our criteria (Figure S5).

### Region of Interest Definition

Based on the topographic patterns and cluster correction results, four ROIs were defined for subsequent analyses. Two ROIs corresponded to confirmatory cluster-corrected effects: anterior fast spindle duration, averaged over a 47-channel anterior ROI capturing the full extent of the effect (whose 29-channel core survived permutation correction at corrected *p* = 0.017; Figure 3a–c), and anterior slow spindle peak frequency, averaged over its 24-channel surviving cluster (corrected *p* = 0.047; Figure 3d–f). Because the 29-channel corrected fast-duration cluster represented the statistically strongest core of a broader, spatially coherent anterior effect, behavioral models averaged fast spindle duration over the full anterior effect extent to improve the reliability of each participant’s ROI estimate; a sensitivity analysis restricting this ROI to only the 29-channel corrected cluster yielded the same pattern of attentional associations (d-prime, omission errors, and commission errors all remained significant with unchanged direction; Table S5). Two additional ROIs were exploratory patterns from uncorrected topographic maps (Figure 4): posterior slow spindle amplitude (40 channels) and anterior slow spindle duration (22 channels); neither survived cluster correction. Channel identifiers for each ROI are listed in Table S1, and full cluster-based permutation statistics for the spindle metrics tested are provided in Table S2. ROI averages for each participant were extracted for behavioral analyses. All four ROIs were tested as predictors of attentional performance, but interpretive emphasis was placed on effects that survived correction and showed convergent behavioral relevance.

### Spindle Features and Attentional Performance

We next examined whether spindle ROI metrics predicted attentional performance on the TOVA using linear regression models adjusted for age, sex, and HI (*N* = 56; Figure 5; full model statistics are reported in Table S3). Anterior fast spindle duration is presented as the primary predictor, with anterior slow spindle peak frequency (the second cluster-corrected effect), anterior slow spindle duration, and posterior slow spindle amplitude reported for comparison. Rather than treating these models as an exploratory screen across all spindle features and TOVA outcomes, we assigned interpretive priority a priori to spindle features meeting two independent criteria: survival of cluster-based permutation correction in the topographic HI–spindle analysis and association with multiple TOVA outcomes in covariate-adjusted models. Convergence on the same feature across these otherwise independent analyses is unlikely to arise from selectively reporting one of many tested combinations.

**Figure 5.**
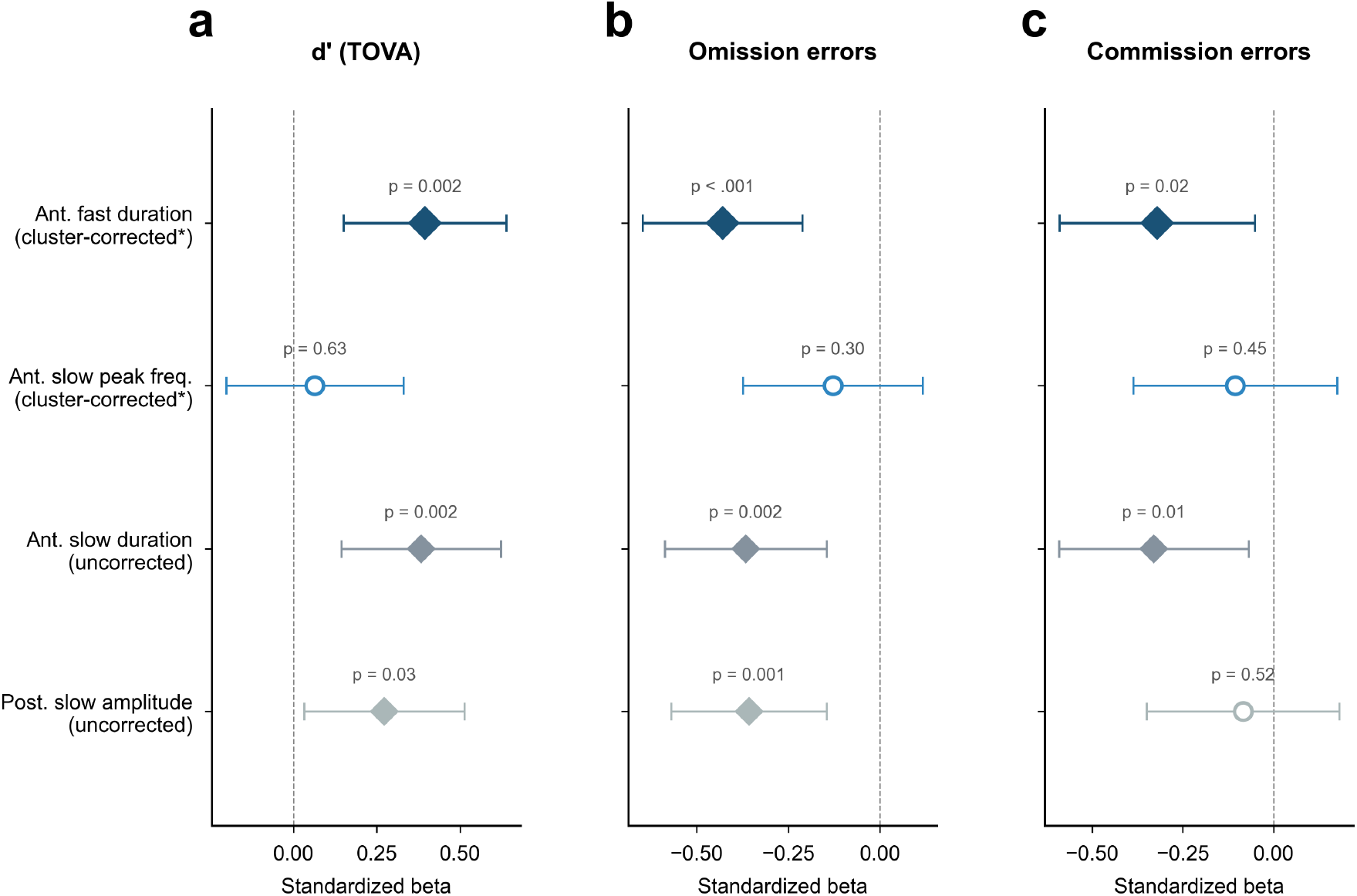
Associations between spindle ROI metrics and attentional performance (TOVA). Standardized regression coefficients with 95% confidence intervals are shown from linear models predicting d-prime (a), omission errors (b), and commission errors (c), adjusted for age, sex, and HI (N = 56). Rows represent spindle metrics derived from data-driven ROIs: anterior fast spindle duration (cluster-corrected p = 0.017; Figure 3), anterior slow spindle peak frequency (cluster-corrected p = 0.047; Figure 3), anterior slow spindle duration (uncorrected; Figure 4), and posterior slow spindle amplitude (uncorrected; Figure 4). Filled diamonds indicate significant predictors (p < 0.05); open circles indicate non-significant associations. Anterior fast spindle duration was both the most robust cluster-corrected topographic effect and the most consistent predictor of attentional performance, whereas the other cluster-corrected effect — slow spindle peak frequency — showed no behavioral associations.

### d-Prime

Greater anterior fast spindle duration was a robust predictor of higher d-prime (β = 0.39, SE = 0.13, *p* = 0.003; Figure 5a), accounting for substantial variance in the model (*R*^2^ = 0.34, *F*(4,51) = 6.58, *p* < 0.001). Subsequent models showed comparable associations: anterior slow spindle duration (β = 0.37, SE = 0.12, *p* = 0.003; *R*^2^ = 0.34) and posterior slow spindle amplitude (β = 0.32, SE = 0.12, *p* = 0.010; *R*^2^ = 0.31). Age was positively associated with d-prime across all three models (β = 0.21–0.27, *p* ≤ 0.011).

### Omission Errors

Anterior fast spindle duration was a significant predictor of fewer omission errors (β = −0.45, SE = 0.11, *p* < 0.001; Figure 5b; *R*^2^ = 0.48, *F*(4,51) = 11.76, *p* < 0.001). Parallel models showed consistent associations for the other ROIs: anterior slow spindle duration (β = −0.38, SE = 0.11, *p* = 0.002; *R*^2^ = 0.44) and posterior slow spindle amplitude (β = −0.39, SE = 0.11, *p* < 0.001; *R*^2^ = 0.45). Age was a robust predictor of fewer omissions across all models (β = −0.24 to −0.31, *p* < 0.01).

### Commission Errors

Greater anterior fast spindle duration (β = −0.28, SE = 0.14, *p* = 0.043; Figure 5c; *R*^2^ = 0.22) and anterior slow spindle duration both predicted fewer commission errors (β = −0.29, SE = 0.13, *p* = 0.028; *R*^2^ = 0.24). Posterior slow spindle amplitude was not a significant predictor of commission errors (β = −0.14, *p* = 0.27), representing the only TOVA outcome where the secondary ROIs diverged from the primary finding. In the primary anterior fast duration model, neither age nor HI was a significant predictor of commission errors.

### Reaction Time

Reaction time was not consistently associated with spindle ROI metrics. None of the four spindle predictors reached significance (all *p* > 0.15). Instead, reaction time was predominantly predicted by age (β = −0.23 to −0.26, *p* < 0.01) and HI (*p* = 0.006–0.03); these ranges span the four ROI models, each of which included age and the hypopnea index as covariates.

## Discussion

This study identifies anterior fast spindle duration as a neurophysiological link between sleep-related respiratory disruption and attentional vulnerability in children. The key finding is not that pediatric SDB broadly reduces spindle activity, but that increasing hypopnea burden is associated with a regionally specific shortening of fast spindle events, and that this same feature is the most consistent spindle predictor of attentional deficits. Thus, the functionally relevant spindle abnormality appears to be a loss of temporal stability within anterior fast thalamocortical oscillations, rather than a generalized failure to generate spindles.

### Regionally Specific Spindle Alterations

Higher hypopnea burden was associated with a spatially structured set of spindle changes rather than a uniform shift across the scalp. Shortened anterior fast spindle duration and slower anterior slow spindle peak frequency survived cluster-based permutation correction, whereas increased posterior slow spindle amplitude and shortened anterior slow spindle duration emerged only in the uncorrected topographic maps. The cluster-corrected fast-duration and slow-frequency effects are taken up in the sections that follow; here we first consider the exploratory slow-spindle effects. The posterior slow spindle amplitude effect, while showing a consistent uncorrected spatial pattern in the primary whole-sample HI model, did not survive cluster-based permutation correction and should be interpreted cautiously. In canonical spindle topography, slow spindles predominate over frontal regions, reflecting engagement of prefrontal thalamocortical circuits [8,9,10]. The HI-associated increase in posterior slow spindle amplitude may reflect disruption of this normal anterior dominance, potentially through fragmentation-induced alterations in thalamocortical gating [6,7,22]. More broadly, the posterior slow amplitude and anterior slow duration effects are best regarded as hypothesis-generating: although they did not survive cluster correction, their convergent associations with attentional performance suggest that they may capture related aspects of spindle morphology. Larger independent samples will be needed to determine whether these exploratory effects replicate and whether developmental stage reliably moderates respiratory effects on slow spindle amplitude.

Follow-up Age × HI analyses support a cautious interpretation of spindle topography. Using a spatially valid permutation null (one shared subject reordering per permutation across all channels, with a channel-forming |*t*| > 2.0 applied identically to the observed and null maps), the primary spindle metrics were largely free of age-moderated HI associations: density, count, and both duration metrics showed no age-moderated effect surviving cluster correction, and only slow spindle amplitude reached corrected significance (cluster *p* = 0.047), therefore we interpret this developmental pattern cautiously rather than as a robust interaction (Figure S2). A cluster-corrected Age × HI interaction was also present for fast spindle peak frequency, whereas the primary anterior fast spindle duration effect itself was not age-moderated. At the level of the predefined anterior duration regions, neither fast nor slow spindle duration showed a significant Age × HI interaction (Table S4, Figure S3), indicating that the HI-related shortening of anterior spindle duration was broadly stable across the 3–11-year range studied here. We caution, however, that the absence of a detectable interaction does not exclude developmental modulation: our cohort spans a relatively narrow, prepubertal window during which frontal thalamocortical circuits are still maturing, and the modest sample size affords limited power to resolve age-by-exposure interactions. Canonical spindle organization was therefore detectable and, on present evidence, broadly stable across this age range, though we cannot consider it developmentally inert.

### Spindle Duration and Attention

The finding that fast spindle duration was the only cluster-corrected HI-sensitive spindle metric to predict attentional outcomes carries important implications for how we conceptualize the neurophysiological impact of SDB on sleep microarchitecture. We use “temporal stability” here to refer to the ability of a spindle event, once initiated, to persist across its oscillatory envelope rather than terminating prematurely. Duration likely reflects this stability of thalamocortical oscillatory bouts, which is thought to depend on reciprocal inhibitory dynamics within the thalamic reticular nucleus and the fidelity of cortical feedback loops [6,22]. One possibility is that respiratory-related arousals truncate spindle events before they complete their full oscillatory envelope; if so, shorter spindle durations would reduce the temporal window available for the synaptic plasticity processes that spindles are thought to support [38,39]. The observation that fast spindles were specifically affected is consistent with evidence that fast spindles engage more extensive cortical networks, including hippocampal-cortical pathways [40,41,42]. The anterior topography of this effect suggests that frontal thalamocortical circuits are preferentially disrupted, consistent with the known vulnerability of prefrontal-dependent processes to sleep disruption in children [3,5].

Two convergent properties of frontal cortex offer a candidate explanation for why these duration effects localized anteriorly rather than over centro-parietal regions, which we advance as a hypothesis rather than a conclusion the present design can establish. First, the prefrontal cortex is the last cortical region to mature: across the first two decades the maximum of sleep slow-wave activity migrates from posterior to frontal cortex in step with regional gray-matter and synaptic maturation, a posteroanterior progression that is still underway across the prepubertal range sampled here [18,19]. The anterior generators sampled in our 3–11-year cohort are therefore among the least synaptically consolidated, and plausibly the least able to sustain stable thalamocortical oscillation when perturbed. Second, and independently, the prefrontal cortex is the region most sensitive to the intermittent hypoxia and sleep fragmentation of pediatric SDB, showing both neurocognitive and neuroimaging evidence of selective injury [1,3,5]. We propose, as a hypothesis for direct test, that these two vulnerabilities—developmental immaturity and respiratory susceptibility—compound, such that a still-maturing frontal circuit has the least reserve to buffer respiratory perturbation and preferentially expresses shortened spindle duration. Because our cohort lies entirely within this prepubertal, maturation-sensitive window, the present cross-sectional design cannot isolate the developmental contribution, and we did not observe an age-by-HI interaction within this range (Table S4); adjudicating this hypothesis will require samples spanning a broader age range and longitudinal designs (see Limitations and Future Directions).

The convergence between topographic and behavioral findings provides the central contribution of this study. Anterior fast spindle duration was not only the most robust cluster-corrected topographic effect of HI but also a consistent predictor of d-prime, omission errors, and commission errors — three TOVA metrics indexing distinct components of attentional functioning. This dual convergence — surviving multiple comparison correction in the topographic domain and showing the robust behavioral associations — argues against a multiple comparison artifact and supports fast spindle duration as a genuine neurophysiological marker. The association with d-prime (discriminatory attention) suggests that spindle duration supports neural processes underlying perceptual discrimination, while associations with omission errors (sustained attention/vigilance) and commission errors (inhibitory control) indicate broader relevance to executive attention. Critically, these associations remained significant after adjusting for age, sex, and HI, indicating that spindle duration is associated with attentional performance independently of respiratory disruption severity and developmental maturation. The absence of significant spindle-attention associations for reaction time — which was instead predicted by age and HI — suggests that anterior fast spindle duration is not merely a marker of generalized psychomotor speed. Instead, its associations point toward accuracy-related attentional control.

### Slow Spindle Peak Frequency

The finding that higher HI was associated with slower slow spindle peak frequency represents the second cluster-corrected effect in this study. Spindle peak frequency is among the most stable, trait-like features of sleep microarchitecture [14] and undergoes systematic developmental changes across childhood and adolescence [20,21], suggesting that it reflects enduring properties of thalamocortical circuit organization. A reduction in oscillation frequency could reflect altered thalamocortical pacing, potentially through hypoxemia-related effects on thalamic reticular nucleus or changes in cortical feedback timing [6,22]. Notably, despite surviving cluster-based correction at the topographic level, anterior slow spindle peak frequency did not predict attentional performance on any TOVA metric. This dissociation is informative: slow spindle peak frequency appears to index respiratory sensitivity of thalamocortical physiology rather than the attention-relevant pathway captured in this study. More broadly, respiratory disruption appears to affect separable dimensions of thalamocortical function — event stability versus oscillatory pacing — with attentional relevance concentrated in spindle duration. This parallels observations that spindle frequency and duration load on separable dimensions of spindle phenotype with distinct trait-like properties [14] and suggests these dimensions may relate to different downstream functional pathways.

### Limitations and Future Directions

Several limitations also define the next steps. The cross-sectional design cannot determine whether respiratory disruption shortens fast spindles, whether altered thalamocortical maturation increases vulnerability to respiratory events, or whether both reflect shared developmental factors. Longitudinal studies before and after SDB treatment provide the clearest test of directionality: if fast spindle duration normalizes with improved breathing and tracks attentional gains, it would strengthen the mechanistic interpretation; if respiratory indices improve without spindle recovery, the spindle phenotype may instead mark a more persistent neurodevelopmental vulnerability. Pediatric treatment studies show that adenotonsillectomy reliably improves respiratory and behavioral outcomes in children with SDB [1], whereas effects on cognition are inconsistent: the largest randomized trial found no significant neurocognitive benefit despite respiratory improvement [43], a controlled study found neurocognitive deficits that persisted after surgery [44], and other reports describe cognitive gains [45]. Extending that framework to hdEEG would clarify whether spindle alterations are reversible and whether recovery of spindle duration corresponds to cognitive improvement. An explicit test of the developmental-vulnerability hypothesis advanced above would require sampling across a wider age range—ideally spanning the pre-to post-pubertal transition—and relating the HI sensitivity of anterior spindle duration to individual markers of frontal maturation, such as the frontal-to-occipital slow-wave-activity gradient [18] or spindle–slow-oscillation coupling [39]. A steeper HI–duration association in children with less mature anterior circuits would support compounded vulnerability, whereas an exposure effect independent of maturation would favor direct respiratory susceptibility.

The behavioral analysis intentionally focused on TOVA Quarter 1, the initial low-target-frequency condition [28], because it provides a constrained test of vigilance under sparse target demands. This choice reduced heterogeneity across the full 21.6-minute task but narrows interpretation. Future analyses should test whether spindle duration predicts performance decay across quarters, differentiates low-from high-target-frequency phases, or explains overnight change when both PM and AM TOVA assessments are available. Those models would separate baseline attentional capacity from sleep-dependent change more directly than a Q1-only cross-sectional model.

The hdEEG approach provided spatial resolution, but these analyses remained scalp-level and combined N2 and N3 sleep. Source localization could identify the thalamocortical generators affected by respiratory disruption, while stage-specific models and spindle-slow oscillation coupling analyses could test whether respiratory burden alters not only spindle morphology but also the timing of spindles within broader NREM coordination [38,39].

## Conclusion

Together, these findings position spindle duration as a measurable sleep-microarchitectural feature associated with attentional vulnerability in childhood respiratory disruption. Conventional respiratory indices summarize airway physiology, but they do not specify how sleep-dependent brain rhythms are altered or why children with similar respiratory burden may differ cognitively. By combining hdEEG topography with TOVA performance, this study identifies shortened anterior spindle duration as a candidate physiological readout of that missing link: a regionally specific weakening of sustained thalamocortical oscillations with functional relevance for attention. The broader implication is that pediatric SDB may be understood not only as disturbed breathing during sleep, but as a disruption of the developing sleep physiology that supports daytime cognitive control.

## Acknowledgments

The authors thank Poorang Nori of Wisconsin Sleep for assistance with the polysomnography setup.

## Code and Data Availability

Analytic code supporting the spindle topographic, cluster-permutation, and behavioral analyses reported here is publicly available at https://github.com/idossha/Pediatric_Spindles_HI. Deidentified participant-level analytic data, together with a data dictionary, may be made available to qualified investigators upon reasonable request after publication; requests should be directed to the corresponding author and will be reviewed by the study investigators and the University of Wisconsin–Madison in accordance with institutional policies, participant consent, and applicable IRB and data-use requirements, under a signed data-use agreement. Raw high-density EEG and polysomnography recordings will not be made publicly available because of their size and complexity and the risk of participant re-identification in a pediatric clinical sample.

## Author Contributions

Conceptualization: SGJ, CM. Data curation: TT, ES. Formal analysis: IH, TT. Funding acquisition: SGJ. Investigation: SGJ, CM, ES, BP, TK, AM, BR. Methodology: SGJ, CM, ES, AMV, IH, TT. Project administration: SGJ, ES, AMV. Resources: SGJ, CM. Software: IH, TT. Supervision: SGJ. Validation: SGJ, IH, TT. Visualization: IH, TT. Writing – original draft: IH, TT, SGJ. Writing – review and editing: IH, SGJ, CM, TT, ES, AMV, BP, TK, AM, BR. Author initials: IH, Ido Haber; TT, Tamara P. Taporoski; BP, Beth Peterson; CM, Camilla Matthews; TK, Tony Kille; AM, Annika Myers; BR, Brady Riedner; ES, Emma Strainis; AMV, Ana Maria Vascan; SGJ, Stephanie G. Jones.

