## Supplementary material for "Sleep-Related Respiratory Disruption is Associated with Altered Spindle Morphology and Poorer Attention in Children": SI

**Table S1.** *Channel membership of the four spindle regions of interest.*

| Region of interest | Band | Metric | *n* | Channel labels |
|---|---|---|---|---|
| Anterior fast duration | Fast | Duration | 47 | 4, 5, 6, 7, 8, 9, 12, 13, 14, 15, 16, 19, 20, 22, 24, 27, 28, 29, 30, 33, 34, 35, 36, 38, 39, 40, 41, 42, 43, 44, 47, 50, 51, 52, 53, 58, 59, 60, 185, 186, 196, 197, 198, 206, 207, 212, 224 |
| Anterior slow peak frequency | Slow | Peak frequency | 24 | 3, 4, 11, 12, 13, 19, 20, 21, 26, 27, 28, 32, 33, 34, 38, 47, 54, 61, 205, 213, 214, 222, 223, 224 |
| Posterior slow amplitude | Slow | Amplitude | 40 | 69, 71, 72, 74, 75, 76, 77, 78, 79, 80, 84, 85, 86, 87, 88, 89, 97, 99, 100, 110, 129, 130, 141, 142, 143, 153, 154, 162, 163, 164, 170, 171, 172, 179, 180, 181, 182, 191, 192, 193 |
| Anterior slow duration | Slow | Duration | 22 | 8, 13, 14, 15, 16, 17, 20, 21, 22, 24, 40, 44, 50, 51, 52, 57, 58, 59, 197, 198, 207, 215 |

*Note. Anterior fast duration and anterior slow peak frequency are the cluster-corrected effects; posterior slow amplitude and anterior slow duration are exploratory regions from the uncorrected topographic maps. The channels listed are the behavioral averaging ROIs: for anterior slow peak frequency this ROI is exactly the 24-channel surviving cluster, whereas for anterior fast duration it spans the full anterior extent of the effect (47 channels) and is broader than the 29-channel surviving cluster (Table S2). Channel labels follow the 172-channel high-density montage (Figure S1).*

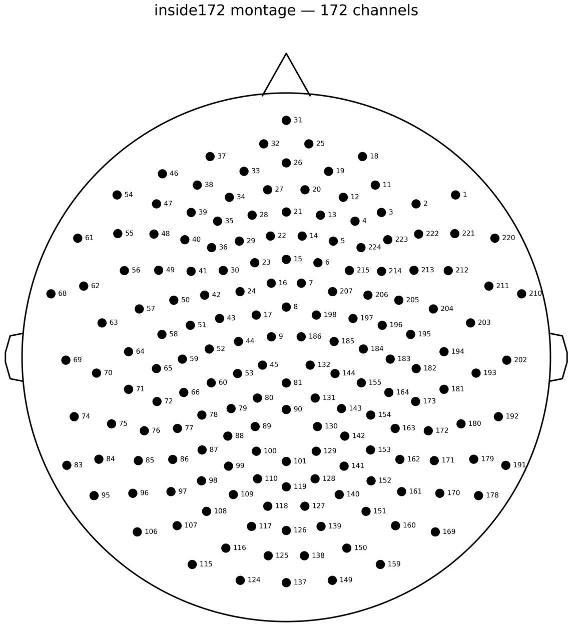


**Figure S1.** *Layout of the 172-channel high-density EEG montage (inside172) used throughout the analysis. Each marker denotes a recording channel; labels are the channel identifiers referenced in Table S1. No data are plotted.*

**Table S2.** *Cluster-based permutation results for hypopnea-related spindle effects.*

| Spindle metric | Cluster size (ch) | Cluster mass (Σ\|*t*\|) | Peak *t* | Corrected *p* |
|---|---|---|---|---|
| Fast duration | 29 | 66.9 | -2.86 | **0.017** |
| Slow peak frequency | 24 | 56.0 | -2.74 | **0.047** |
| Slow amplitude | 15 | 33.9 | +2.64 | 0.087 |
| Slow duration | 4 | 8.9 | -2.43 | 0.279 |
| Slow density | 0 | — | — | — |
| Fast density | 1 | 2.1 | -2.06 | 0.363 |

*Note. Association between the hypopnea index and each spindle metric (channel-wise model adjusted for age and sex; cluster-forming $|t| > 2.0$; 5,000 Freedman–Lane residual permutations). Corrected p is the proportion of permutations whose maximal cluster mass met or exceeded the observed value; significant values (< 0.05) are shown in bold. Em dashes denote metrics with no suprathreshold cluster.*

**Table S3.** *ROI predictors of attentional performance.*

| TOVA outcome | ROI predictor | *β* | SE | *t* | *p* | 95% CI | Adj. $R^2$ |
|---|---|---|---|---|---|---|---|
| d-prime | Anterior fast duration | +0.387 | 0.125 | +3.09 | **0.003** | [0.14, 0.64] | 0.289 |
| d-prime | Anterior slow peak frequency | +0.035 | 0.136 | +0.26 | 0.797 | [-0.24, 0.31] | 0.157 |
| d-prime | Anterior slow duration | +0.372 | 0.121 | +3.08 | **0.003** | [0.13, 0.61] | 0.288 |
| d-prime | Posterior slow amplitude | +0.320 | 0.120 | +2.67 | **0.010** | [0.08, 0.56] | 0.259 |
| Omission errors | Anterior fast duration | -0.449 | 0.111 | -4.03 | **0.000** | [-0.67, -0.23] | 0.439 |
| Omission errors | Anterior slow peak frequency | -0.127 | 0.126 | -1.01 | 0.319 | [-0.38, 0.13] | 0.275 |
| Omission errors | Anterior slow duration | -0.375 | 0.112 | -3.37 | **0.002** | [-0.60, -0.15] | 0.395 |
| Omission errors | Posterior slow amplitude | -0.387 | 0.107 | -3.63 | **0.001** | [-0.60, -0.17] | 0.412 |
| Commission errors | Anterior fast duration | -0.282 | 0.136 | -2.07 | **0.043** | [-0.56, -0.01] | 0.163 |
| Commission errors | Anterior slow peak frequency | -0.056 | 0.140 | -0.40 | 0.689 | [-0.34, 0.23] | 0.095 |
| Commission errors | Anterior slow duration | -0.295 | 0.130 | -2.27 | **0.028** | [-0.56, -0.03] | 0.176 |
| Commission errors | Posterior slow amplitude | -0.145 | 0.131 | -1.10 | 0.275 | [-0.41, 0.12] | 0.114 |
| Reaction time | Anterior fast duration | -0.175 | 0.123 | -1.43 | 0.160 | [-0.42, 0.07] | 0.318 |
| Reaction time | Anterior slow peak frequency | -0.009 | 0.124 | -0.08 | 0.939 | [-0.26, 0.24] | 0.291 |
| Reaction time | Anterior slow duration | -0.115 | 0.120 | -0.96 | 0.339 | [-0.36, 0.12] | 0.303 |
| Reaction time | Posterior slow amplitude | -0.129 | 0.116 | -1.11 | 0.272 | [-0.36, 0.10] | 0.307 |

*Note. Standardized linear-regression coefficients for each ROI predicting each TOVA attention outcome, adjusted for age, sex, and the (log) hypopnea index ($N = 56$). Coefficients are in standard-deviation units; the adjusted $R^2$ is for the full model. Significant p-values (< 0.05) are shown in bold.*

### Age × Hypopnea Index Interaction

To test whether the whole-sample hypopnea-index (HI) associations masked developmentally distinct effects, each spindle metric was refit channel-wise with an Age × log(HI + 1) interaction term, adjusting for age, sex, and the HI main effect (mean-centered predictors) in the full analytic sample (N = 62). The interaction term was evaluated with a Freedman–Lane residual permutation test (5,000 permutations; channel-forming threshold of $|t| > 2.0$—the two-sided $p < .05$ critical value, applied identically to the observed and permutation-null t-maps and matching the main HI cluster test) with a spatially valid cluster correction over the channel adjacency, in which a single shared subject reordering is applied across all channels per permutation (permuting channels independently makes the null spatially white and inflates significance). Under this corrected procedure, among the primary metrics—density, count, duration, and amplitude—only slow spindle amplitude showed an age-moderated HI association that reached corrected significance, and then only narrowly: it formed a 25-channel anterior cluster (cluster $p = 0.047$, mean $t = -2.32$). Because this p-value sits just below the 0.05 threshold and the underlying slow-amplitude HI effect is itself exploratory (it did not survive correction in the whole-sample analysis), we treat this developmental pattern cautiously rather than as a robust interaction (Figure S2): the association between HI and slow spindle amplitude was positive in younger children and flattened to negative in older children. Density, count, and both duration metrics showed no age-moderated HI association surviving correction. Among secondary metrics, fast spindle peak frequency showed a cluster-corrected interaction (33-channel cluster, $p = 0.012$); this metric is not part of the primary duration-focused analysis and is documented for completeness. Crucially, the primary anterior fast-duration effect was not moderated by age and remained stable in the whole-sample model.

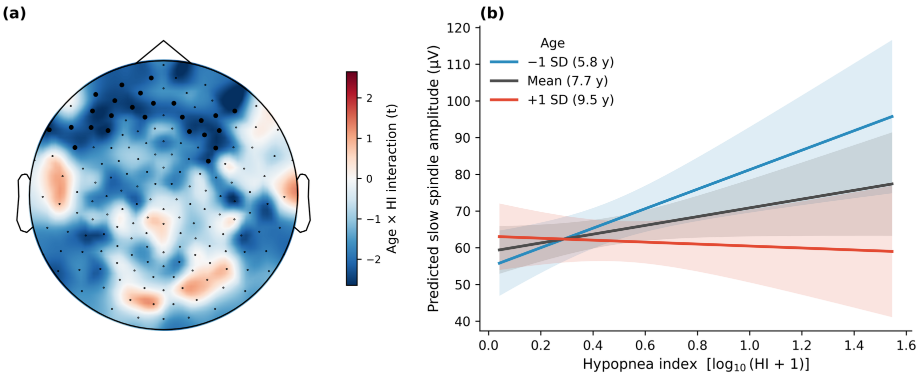


**Figure S2.** *Exploratory Age × hypopnea-index interaction for slow spindle amplitude (N = 62). (a) Channel-wise Age × log(HI + 1) interaction t-map; black markers denote the largest (25-channel) anterior cluster, which narrowly reached corrected significance under the spatially valid cluster-based permutation correction (cluster p = 0.047). (b) Model-estimated slow spindle amplitude as a function of hypopnea index at the mean age and ±1 SD (−1 SD, Mean, +1 SD; adjusted for sex), averaged over the cluster region; shaded bands are 95% confidence intervals. The association between hypopnea index and slow spindle amplitude is positive in younger children and flat-to-negative in older children; because this developmental pattern sits just past the 0.05 threshold and the underlying slow-amplitude HI effect is itself exploratory, it should be interpreted cautiously and regarded as hypothesis-generating.*

To test the same question within the predefined regions of interest—rather than across the whole channel array—each ROI series (Table S1) was refit with an Age × log(HI + 1) interaction term (adjusting for age, sex, and the HI main effect; N = 62). Because each ROI is a single averaged series, this is a single ordinary least-squares interaction model per ROI and does not depend on the cluster-null assumptions of the channel-wise test. Simple slopes were probed at the mean age and ±1 SD (Aiken and West) rather than at arbitrary ages, so that every probed age falls within well-sampled data. Neither the anterior fast-duration ROI (interaction p = 0.77) nor the anterior slow-duration ROI (p = 0.44) showed a significant Age × HI interaction (Table S4, Figure S3): the negative association between hypopnea index and anterior spindle duration was broadly stable across the sampled age range. This stability is consistent with—but does not by itself establish—a developmentally invariant respiratory effect on duration; as noted in the main text, the narrow prepubertal age window and modest sample size limit power to detect age-by-exposure interactions, so this null is not evidence against developmental modulation.

**Table S4.** *ROI-level Age × hypopnea-index interaction for the spindle regions of interest.*

| Region of interest | Age × HI $\beta$ | $t$ | $p$ | HI slope (−1 SD) | HI slope (Mean) | HI slope (+1 SD) |
|---|---|---|---|---|---|---|
| Anterior fast spindle duration | +0.0055 | +0.29 | 0.773 | -0.109 | -0.099 | -0.089 |
| Anterior slow spindle duration | -0.0132 | -0.77 | 0.443 | -0.053 | -0.077 | -0.101 |
| Posterior slow spindle amplitude | -3.2108 | -1.52 | 0.134 | +16.656 | +10.814 | +4.973 |
| Anterior slow spindle peak frequency | +0.0684 | +1.94 | 0.057 | -0.317 | -0.193 | -0.068 |

*Note. Each predefined ROI series (Table S1) was fit with roi ~ age_c + logHI_c + age_c:logHI_c + sex (N = 62, mean-centered predictors). The interaction coefficient (β, t, p) tests whether age moderates the hypopnea-index association; simple slopes give the model-implied association between log(HI + 1) and the ROI metric probed at the mean age and ±1 SD (mean = 7.7, SD = 1.8; i.e. 5.9, 7.7, and 9.5 years), following Aiken and West. Units per ROI: duration in s, amplitude in µV, peak frequency in Hz. Neither anterior duration ROI shows a significant Age × HI interaction, indicating a developmentally stable hypopnea effect across the 3–11-year range. Significant p-values (< 0.05) are shown in bold.*

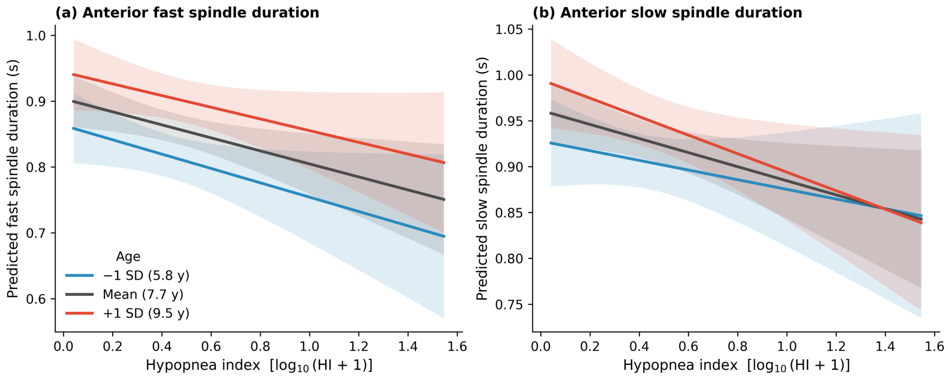


**Figure S3.** *ROI-level Age × hypopnea-index interaction for the two anterior spindle-duration regions (N = 62). Model-estimated spindle duration as a function of hypopnea index at the mean age and ±1 SD (−1 SD, Mean, +1 SD; adjusted for sex), for (a) the anterior fast-duration ROI and (b) the anterior slow-duration ROI; shaded bands are 95% confidence intervals. The HI–duration slopes are negative and broadly parallel across ages, and the Age × HI interaction is non-significant for both ROIs (Table S4), indicating that the hypopnea-related shortening of anterior spindle duration does not vary detectably with age within this prepubertal sample.*

### Hypopnea index distribution and robustness of the primary effect

The hypopnea index was strongly right-skewed (skewness = 4.49; range 0.1–34.1 events/hour), so all regression models used log10(HI + 1), which substantially reduced the skew (1.10) (Figure S4a–b). To confirm that the primary cluster-corrected effect (shorter anterior fast spindle duration with rising HI; cluster p = 0.017) was not driven by any single participant, the cluster-based permutation test was refit 62 times, each omitting one child (leave-one-subject-out). The corrected cluster p remained below .05 in 47 of 62 folds (leave-one-out p range 0.005–0.131; Figure S4c–d). The effect was most sensitive to the single child with the highest hypopnea index (HI = 34.1), whose omission attenuated the cluster (p = 0.131); the remaining folds that exceeded .05 reached at most p = 0.109, consistent with the modest power of a cluster-corrected effect at this sample size. Because counting folds against a fixed significance threshold is itself unstable at this sample size, effect stability was quantified directly with a subject-level bootstrap (10,000 resamples with replacement) of the cluster-averaged, covariate-adjusted standardized association between log(HI + 1) and fast spindle duration. The association was negative in 99.9% of resamples, with a standardized coefficient of −0.31 (95% bootstrap CI [−0.49, −0.12]) that excluded zero (Figure S4d), indicating that the shortening of fast spindle duration with rising hypopnea index is robust to subject composition rather than contingent on individual participants.

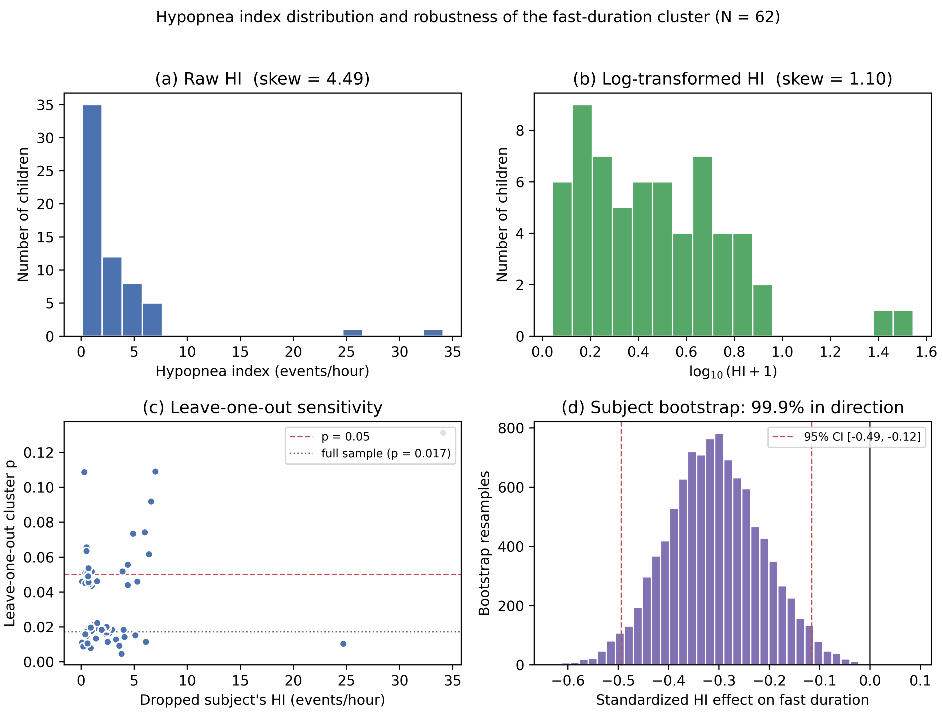


**Figure S4.** *Hypopnea index distribution and robustness of the fast-duration cluster (N = 62). (a) Raw hypopnea index, showing strong right skew. (b) log10(HI + 1), the mean-centered predictor used in all models. (c) Leave-one-subject-out corrected p for the primary fast-duration cluster plotted against the dropped child's hypopnea index; the dashed line marks p = .05 and the dotted line the full-sample value (p = 0.017). (d) Subject-level bootstrap (10,000 resamples) of the cluster-averaged standardized association between log(HI + 1) and fast spindle duration (adjusted for age and sex); dashed lines mark the 95% percentile confidence interval, which excludes zero, and the effect is negative in 99.9% of resamples.*

### Sensitivity of the anterior fast-duration behavioral ROI

The behavioral models averaged anterior fast spindle duration over the full 47-channel anterior extent of the hypopnea-index effect (channels significant at $p < 0.05$ together with contiguous trending channels at $p < 0.08$; Table S1), which is broader than the 29-channel subset that survived cluster-based permutation correction (Table S2). The broader region was used because averaging a per-participant metric over the full spatially coherent effect improves the reliability of each participant's ROI value; cluster correction is a family-wise-error threshold on the topographic test, not a boundary on where the effect is measurable, and the trending channels immediately adjacent to the surviving cluster carry the same negative association. Because the 47-channel ROI is defined from the same hypopnea-index t-map, it is a superset of the corrected cluster and shares its sign. To confirm that this choice does not drive the cognitive findings, every attention model was refit using fast spindle duration averaged over only the 29-channel corrected cluster. The pattern was unchanged (Table S5): higher fast spindle duration predicted higher d-prime and fewer omission and commission errors (all $p < 0.05$), with no association for reaction time, and if anything the effects were marginally stronger for the 29-channel cluster.

**Table S5.** *Sensitivity of the anterior fast-duration behavioral results to ROI definition.*

| TOVA outcome | *β* (47-ch ROI) | *p* (47-ch) | *β* (29-ch cluster) | *p* (29-ch) |
|---|---|---|---|---|
| d-prime | +0.387 | **0.003** | +0.405 | **0.002** |
| Omission errors | -0.449 | **0.000** | -0.459 | **0.000** |
| Commission errors | -0.282 | **0.043** | -0.303 | **0.030** |
| Reaction time | -0.175 | 0.160 | -0.196 | 0.115 |

*Note. Standardized anterior fast spindle duration predicting each TOVA outcome (linear models adjusted for age, sex, and the log hypopnea index; N = 56), computed with the primary 47-channel anterior ROI (the full spatial extent of the effect) versus the 29-channel permutation-corrected cluster (Table S2). The two definitions yield the same pattern of results: d-prime, omission errors, and commission errors are significant with unchanged sign, and reaction time is non-significant, under both definitions. Significant p-values (< 0.05) are shown in bold.*

### Spindle density topography

Neither slow nor fast spindle density showed a robust topographic association with the hypopnea index. In the whole-sample channel-wise model (adjusted for age and sex; N = 62), slow spindle density was weakly positive over posterior channels but reached significance at no individual channel (peak $|t| = 1.97$; all $p > 0.05$, uncorrected), and no suprathreshold cluster formed under permutation correction (cluster p = n/a). Fast spindle density showed a weak, predominantly negative pattern, with a single channel reaching $p < 0.05$ and a largest cluster that did not survive correction (cluster $p = 0.363$).

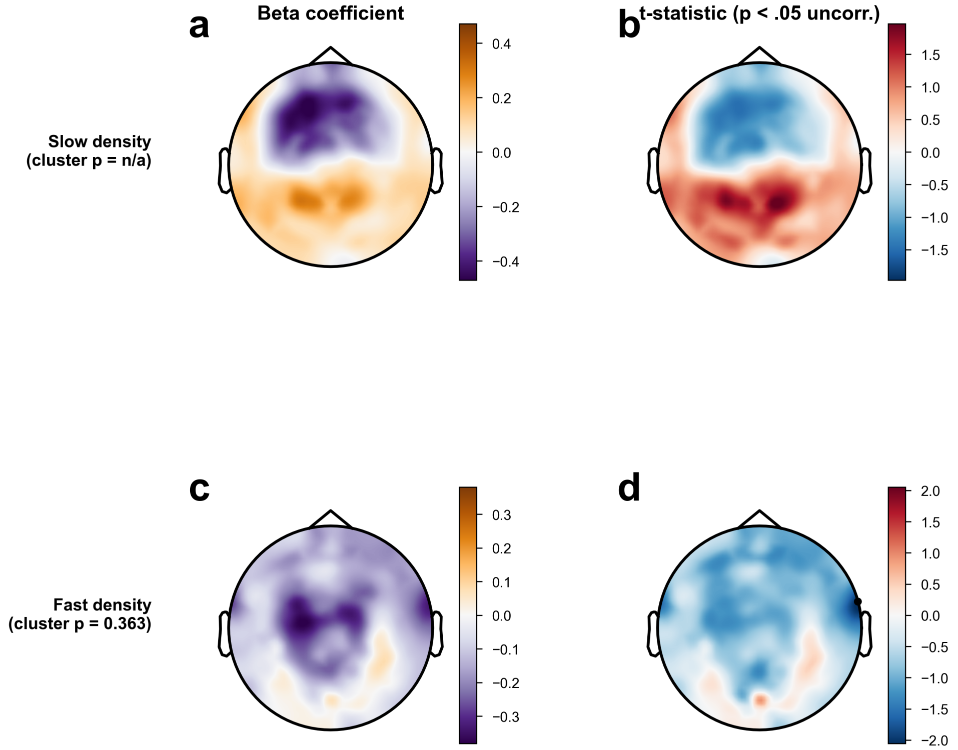


**Figure S5.** *Channel-wise association between the hypopnea index and spindle density. Beta coefficients (left) and t-statistics (right) across the 172-channel montage for (a, b) slow spindle density and (c, d) fast spindle density.*
